# Eye contact in active and passive viewing: event-related brain potential evidence from a combined eye tracking and EEG study

**DOI:** 10.1101/669341

**Authors:** T. Stephani, K. Kirk Driller, O. Dimigen, W. Sommer

## Abstract

Eye contact is a salient social cue which is assumed to influence early brain processes involved in face perception. The N170 component in the event-related potential (ERP) has frequently been reported to be larger to faces with an averted rather than direct gaze towards the observer. In most studies, however, this effect has been investigated in comparatively artificial, passive settings where participants were instructed to fixate their gaze while observing occasional gaze changes in stimulus faces. Yet, it is unclear whether similar mechanisms are in place during naturalistic gaze interactions involving the continuous interplay of directed and averted gaze between the communication partners. To fill this gap, we compared passive viewing of gaze change sequences with an active condition where participants’ own gaze continuously interacted with the gaze of a stimulus face; while recording ERPs and monitoring gaze with eye tracking. In addition, we investigated the relevance of emotional facial expressions for gaze processing. For both passive viewing and active interaction, N170 amplitudes were larger when the gaze of stimulus faces was averted rather than directed at the participants. Furthermore, eye contact decreased P300 amplitudes in both conditions. Emotional facial expression influenced N170 amplitudes but did not elicit an early posterior negativity nor did it interact with gaze direction. We conclude that comparable mechanisms of gaze perception are in place in gaze interaction as compared to passive viewing, encouraging the further study of the eye contact effect in naturalistic settings.

## 1 Introduction

Eye gaze behavior plays a crucial role in human social interactions (Hamilton, 2016): Another person’s gaze provides visual feedback, helps to regulate the flow of conversation, communicates emotions, and provides information about the communication partner’s attention (Kendon, 1967; Vertegaal, Slagter, van der Veer, & Nijholt, 2001). During face-to-face communication, the partners typically move their gaze back and forth between prominent areas of the other person’s face, such as the eye and mouth region (Lusk & Mitchel, 2016), and frequently make and break eye contact.

Many studies indicate that both *eye contact* – when two persons mutually look at each other’s eyes – as well as *direct gaze* – when someone is looked at regardless of eye contact – have a special impact on human cognition, since we tend to deploy visual attention to faces looking at us (Conty, Coelho, Tijus, Hugueville, & George, 2006; Lyyra, Astikainen, & Hietanen, 2017; Senju & Hasegawa, 2005). This effect presumably emerges since direct gaze signals an intention of communication (Senju & Johnson, 2009) and cues reciprocal attention. Impairments in eye contact processing can be observed in disorders characterized by aberrant social behavior and communication deficits, such as autism (Pelphrey, Morris, & McCarthy, 2005), schizophrenia (Tso, Mui, Taylor, & Deldin, 2012), and social anxiety (Schneier, Rodebaugh, Blanco, Lewin, & Liebowitz, 2011). It is therefore important to understand the neural mechanisms of processing eye gaze.

However, most studies on the topic have used rather artificial, passive stimulation paradigms, where the gaze of the stimulus face changes independently of the observer’s gaze. Furthermore, studies on face processing have usually used trial-by-trial paradigms, briefly presenting different faces every few seconds, which is at variance with natural communication situations that involve a continuous interplay of gaze direction. The present study therefore aimed to extend eye contact-related findings from passive viewing to an interactive setting with a continuous interplay of eye gaze that more closely resembles natural human communication. Because the perception and interpretation of eye gaze may be influenced by the facial expression of the communication partner, we also investigated the interactions between gaze perception and emotional facial expressions.

### 1.1 Electrophysiological correlates of eye contact

The neural underpinnings of gaze perception have been investigated with fMRI and event-related potentials (ERP). Here we focus on ERPs due to their high temporal resolution.

#### 1.1.1 N170

The most prominent ERP component in face perception research is the N170, a negative deflection peaking at around 170 ms after stimulus onset at lateral occipito-temporal electrodes, typically more pronounced over the right hemisphere. The N170 is thought to reflect the perceptual encoding of facial structures (Eimer, 2000), particularly of eyes (Bentin, Allison, Puce, Perez, & McCarthy, 1996). Structural encoding of the eyes might be crucial when it comes to human social interaction since the fast and reliable perception of gaze cues is essential for successful communication. It is therefore not surprising that the N170 is sensitive to gaze direction. In many studies, larger N170 amplitudes have been found for faces with averted gaze as compared to faces with direct gaze (Bagherzadeh-Azbari, Lion, Dimigen, Stephani, & Sommer, in preparation; Itier, Alain, Kovacevic, & McIntosh, 2007; Latinus et al., 2015; Puce, Smith, & Allison, 2000). Yet, there are also reports about larger N170 amplitudes for direct than averted gaze (Conty, Dezecache, Hugueville, & Grezes, 2012; Conty, N’Diaye, Tijus, & George, 2007). Latinus et al. (2015) suggested that these inconsistencies may be task-related. In their study they compared a spatial and a social task and found larger N170 amplitudes for averted than for direct gaze when the direction of the observed gaze shifts was to be indicated, whereas the effect was attenuated for an eye-contact detection task.

#### 1.1.2 Other ERP components

In addition to the N170 component, eye contact effects have been found in the P300, a classic index of the active attentive evaluation of stimuli (Cuthbert, Schupp, Bradley, Birbaumer, & Lang, 2000; Donchin & Coles, 1988). Specifically, Conty et al. (2007) observed larger amplitudes for eye contact as compared to gaze aversion in both subcomponents of the P300, the P3a and P3b, which may reflect the typically observed behavioral effect of eye contact drawing the observer’s attention. Note, however, that also the eye contact effects on late positive ERP components are not entirely consistent, as a centro-parietal “P350” component reported by Puce et al. (2000) showed larger amplitudes for averted gaze than for eye contact.

#### 1.1.3 Interaction of gaze direction and emotion

Information conveyed by another person’s gaze strongly depends on contextual factors, such as the person’s facial expression, body gesture, and identity (Conty et al., 2012; Hamilton, 2016). It therefore appears conceivable that the perception of gaze direction and emotional facial expression interact. In case of neutral or happy facial expressions, the observer might interpret direct gaze as a cue for intended communication (i.e., approach behavior; Hietanen, Leppanen, Peltola, Linna-Aho, & Ruuhiala, 2008), whereas eye contact with an angry face may be appraised as a threat (Klucharev & Sams, 2004) and trigger fight-or-flight.

A prominent ERP component linked to emotional processing is the early posterior negativity (EPN), a relative negativity at occipito-temporal electrodes, which is more pronounced for emotional than neutral stimuli (Schupp et al., 2004; Schupp, Junghöfer, Weike, & Hamm, 2003). For facial expressions, the effect typically arises around 150 to 200 ms post-stimulus and lasts until around 300 ms (Schacht & Sommer, 2009). The EPN is thought to reflect enhanced sensory encoding resulting from reflex-like visual attention to emotional stimuli (Rellecke, Palazova, Sommer, & Schacht, 2011; Schupp et al., 2004), enabling their subsequent, elaborate processing (Schacht & Sommer, 2009). Furthermore, there is evidence that emotion effects on the EPN can be modulated by face orientation (Bublatzky, Pittig, Schupp, & Alpers, 2017) as well as by gaze direction. In line with this, Bagherzadeh-Azbari et al. (in preparation) found that angry and happy faces induced a larger EPN compared to neutral faces, which, however, only held true for gaze changes from averted to direct but not from direct to averted gaze. Thus, emotional expressions seem to be differentially processed depending on the gaze direction of the observed face.

### 1.2 Extending laboratory studies to more naturalistic situations

Most of the studies above investigated eye contact in settings where participants kept their gaze fixated at the same location of a computer screen and had no control over the stimulus presentation. Thus, participants *passively observed* the stimuli instead of *interacting* with them as during real-life situations. However, some studies have started to bridge the gap between these conventional and rather artificial stimulus presentation paradigms and more naturalistic settings. For example, Pönkänen, Alhoniemi, Leppanen, and Hietanen (2011) found larger N170 amplitudes for direct gaze as compared to averted gaze when participants viewed faces of real persons. Note that this finding is at variance with the studies summarized above, which have studied gaze effects in response to pictures (e.g., Latinus et al., 2015; Puce et al., 2000). Also Myllyneva and Hietanen (2015) presented live human faces while manipulating participants’ belief of being watched. When participants believed to be watched by the stimulus person, a more positive P3 component emerged for direct than for averted gaze. In the present study, we aimed to make two further steps towards more naturalistic gaze interactions.

First, we employed dynamic gaze changes in face stimuli that were continuously present on the screen. In most recent studies, dynamic gaze changes were presented trial-by-trial (e.g., Conty et al., 2012; Latinus et al., 2015), with a fixation cross usually presented first, followed by a face with a certain gaze direction that changed after a short interval. Face identity typically varied from trial to trial, and blank screens were presented in-between trials. Yet in real-life interactions, the face of the partner remains continuously present during the interaction, and communication involves a continuous interplay between eye contact and averted gaze. Although there have been successful approaches to investigate eye contact in continuous-trial designs (Puce et al., 2000), it remains to be determined whether the N170 eye contact effect is a stable phenomenon that can also be observed across longer sequences of gaze change.

As a second step, we investigated the effect of interaction of the participant with the stimulus. It is currently an open question to what extent the results of passive paradigms generalize to real-world social interactions (Schilbach et al., 2013). In fact, the impact of an interactive setting has been demonstrated for various psychological domains: For example, Richardson, Dale, and Kirkham (2007) showed that eye movements of two persons are coupled in dialogues (i.e., interact with each other), and that visual attention is socially coordinated, facilitating successful communication. Active participation as compared to passive observation also facilitates emotional engagement in joint attention situations, which, in turn, recruits brain processes associated with social interaction (Schilbach et al., 2013). On this background, it seems relevant to investigate the effects of eye contact in an interactive as compared to a passive design.

### 1.3 Objectives and hypotheses

In summary, the current study aimed to move the investigation of eye-contact-related ERPs closer towards a real-world setting. We combined eye tracking with concurrent EEG recordings and used a gaze-contingent experimental design to compare eye contact effects in an active viewing paradigm with those found during the more traditional, passive viewing of gaze changes. In the *active viewing* condition, a face presented on the screen changed its gaze direction whenever the participant looked at the eye region of the face, similar to dyadic communication. In the *passive viewing* condition, participants observed gaze changes without interacting with them. In both conditions, gaze changes were presented in a continuous-trial design. That is, rather than seeing a face for only one or two seconds, participants viewed the same face for a longer period of time comprising between 12 to 14 gaze changes.

To maintain the participants’ attention to the stimuli without interfering with the eye contact effect (see Latinus et al., 2015), we used a non-social gaze change counting task. Based on a previous passive viewing study using the same stimulus material (Bagherzadeh-Azbari et al., in preparation), we expected larger N170 amplitudes for gaze aversion, that is, changes from direct to averted gaze, as compared to making eye contact, that is, changes from averted to direct gaze. If the typical N170 eye contact effect generalizes to a more naturalistic setting, this should hold true both for passive and active viewing. But possibly, the effects could be stronger in the active condition since the ability to interact with the stimulus might increase the relevance of picture changes to the participant.

From a methodological view, the most apparent difference between the two viewing conditions was the extensive eye movements in the active condition. As it is well known that every fixation following a saccade goes along with a brain response reflecting the visual input at fixation onset, the so-called fixation-related potential (FRP; Dimigen, Sommer, Hohlfeld, Jacobs, & Kliegl, 2011; Yagi, 1979), we accounted for the overlap of FRPs and subsequent ERPs in the active condition.

In order to investigate possible interactions of emotion and gaze perception, we presented stimulus faces with angry, happy, and neutral facial expressions. We explored whether gaze direction-dependent EPN effects, as observed in a previous study with traditional trial-by-trial presentation and passive viewing (Bagherzadeh-Azbari et al., in preparation), would also arise during continuous active viewing.

## 2 Methods

### 2.1 Participants

Of 23 participants recruited for the study, two had to be excluded from data analyses due to problems with the eye tracking. The average age of the 21 remaining participants (12 female) was *M* = 26.4 years (*SD* = 5.6); all except for one were right-handed (lateralization score, *M* = +95.24, *SD* = 21.82; Oldfield, 1971), and reported normal or corrected-to-normal vision and no psychiatric or neurological disorders. The experiment conformed to the Declaration of Helsinki (2008). Participants provided written informed consent before the study and were reimbursed with course credits.

### 2.2 Stimuli

To create the face stimuli, an initial set of face photographs was taken from the Radboud Faces Database (Langner et al., 2010), which includes pictures with different gaze directions of the same persons and expressions. We selected frontal-view pictures of 14 Caucasian individuals with three emotional expressions: angry, happy, and neutral (IDs of the selected pictures are provided in Supplement 1). To ensure that only eye gaze (but not any other facial features) would change between the seamlessly presented pictures with different gaze directions, we manipulated the stimuli as follows: For each individual and emotional expression, the eye region of the picture with averted gaze was copied and carefully pasted into the eye region of the corresponding picture with direct gaze using Adobe Photoshop software (version CC2015, Adobe Systems, San Jose, CA). When corresponding pictures with different gaze directions were shown in immediate succession, the impression of a dynamic gaze change of the eye region within an otherwise perfectly immobile face resulted (see Fig. 1). Finally, external features like hair, ears, and shoulders were masked, leaving only internal facial features. The set of stimuli is available upon request from the authors.

**Figure 1.**
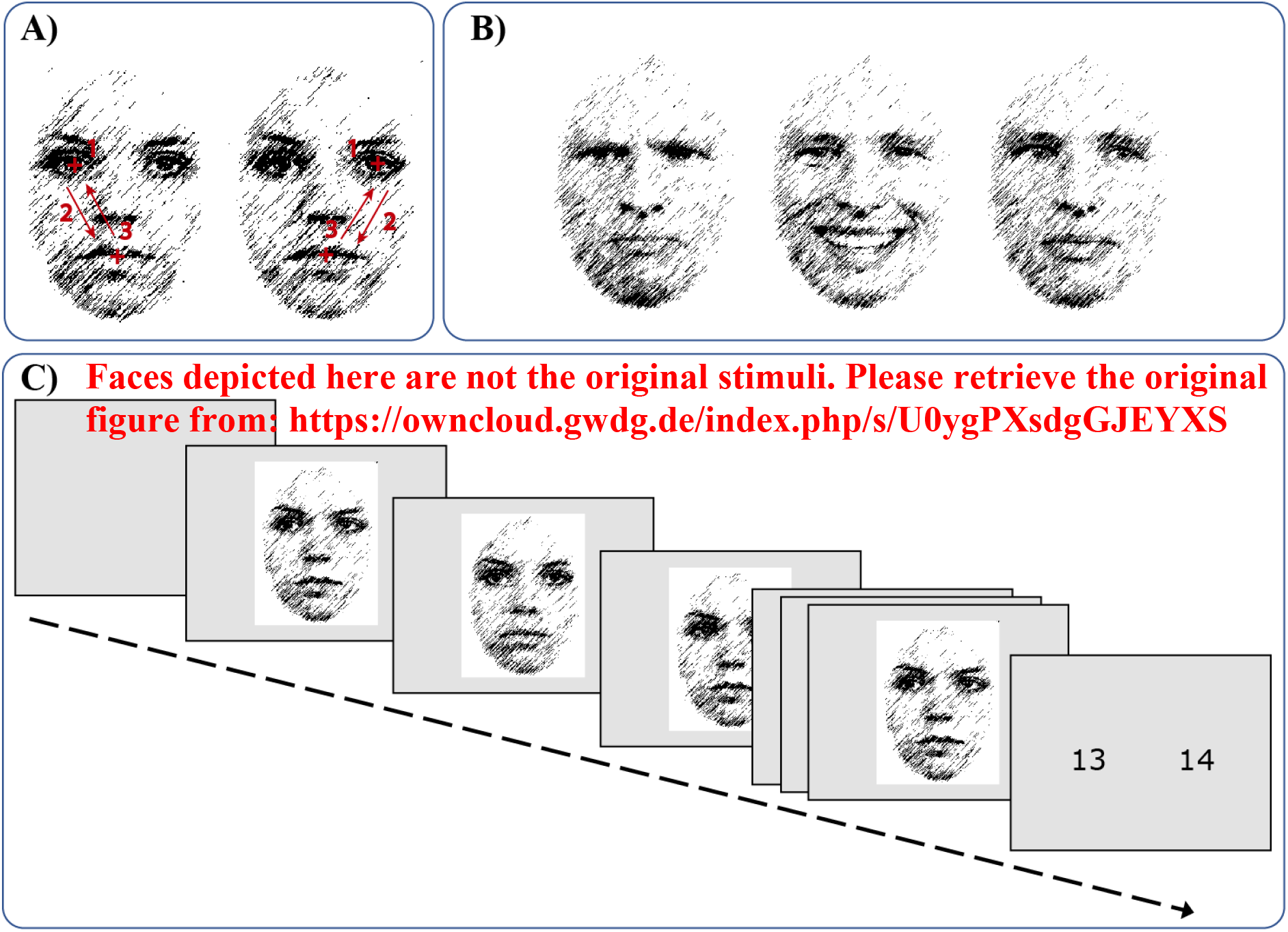
Stimuli and trial structure. A) Instructed gaze movements in the active condition: shift from the mouth to the eye, and back to the mouth; both for the left and right eye. B) Emotional face expressions: angry, happy, and neutral. C) Continuous sequence of gaze changes and subsequent response screen. PLEASE NOTE: Due to bioRxiv’s policy to avoid photographs and any other identifying information of people, cartoon versions of our original stimulus material are shown in this figure. Please retrieve the original figure from: https://owncloud.gwdg.de/index.php/s/U0ygPXsdgGJEYXS

### 2.3 Procedure

After electrode preparation and eye tracker calibration, participants were seated in an electrically shielded cabin with their head position stabilized by the chin rest of the eye tracker. Stimuli were presented at a size of 10 cm × 13.5 cm with a light grey background on a 22-inch CRT monitor (IIyama Vision Master Pro 512, 160 Hz sampling rate, 1024 x 768 pixel). At the given viewing distance of 58 cm, face stimuli covered 9.85° × 13.28° visual angle. Prior to the experiment, prototypical eye movement and blink artifacts were recorded (Dimigen et al., 2011) that were later used in the ocular artifact correction procedure (see section *2.6 Data Analysis*).

The experiment itself was divided into two blocks, the active and the passive viewing condition. In the active viewing condition, participants were instructed to first fixate on the mouth of the stimulus face and then shift their gaze towards the left or right eye, determined by instruction (Fig. 1A). Whenever the participant’s gaze entered the eye region, defined by an invisible rectangular boundary of 2.59 × 2.59°, centered on the midpoint of the pupil, a gaze change of the picture was elicited with a probability of 90% (*change events*; as opposed to 10% *non-change events*). This gaze change occurred 500 ms after the gaze of the participant crossed the boundary during the saccade from the mouth towards the targeted eye of the stimulus face. The interval of 500 ms between eye fixation and stimulus gaze change was considered to approximate an interlocuter’s natural reaction time to direct gaze.

The 10% non-change events served the experimental task, which required counting the overall number of gaze changes in the trial. In addition, the non-change events allowed to compute a clean waveform for the brain activity elicited by the participants saccade towards the eye region, which was un-confounded by the ERP deflections elicited by changes in the stimulus face.

In the 90% of events that contained a stimulus gaze change, six types of gaze changes were possible: left-averted to direct gaze (LD), right-averted to direct (RD), direct to left-averted (DL), direct to right-averted (DR), left-averted to right-averted (LR), and right-averted to left-averted (RL); all directions are given from the participant’s perspective.

The 500-ms interval between eye fixation and gaze change was chosen to separate fixation-related potentials (FRP) elicited by the offset of the saccade to the eye region (see below) from the subsequent stimulus gaze change-related ERPs. In addition, the experimental program required the participant’s gaze to remain inside the eye region for these 500 ms in order to guarantee that the participant was still fixating on the eye when the picture change occurred. In case of an earlier departure from the eye region, no gaze change of the picture was elicited. Participants were instructed to maintain fixation on the eye for about one second and then to look back to the mouth from where they should start a new saccade to one of the eyes (whenever they wanted) eliciting the next gaze change of the face.

In the passive viewing condition, participants fixated on one of the eyes of the face for the whole trial. In this condition, gaze changes of the picture occurred at a randomly jittered interstimulus interval (ISI) between 2000 and 3000 ms (uniform distribution). This ISI approximately corresponded to the average time between two gaze events in the active viewing condition as confirmed by a preceding pilot study with five participants (not reported here).

The order of blocks with active and passive viewing was balanced over participants. In addition, both conditions were counterbalanced for the targeted left or right eye of the stimulus face (resulting in four blocks: active-left, active-right, passive-left, passive-right). In each block, 14 face identities were presented, each with an angry, happy, and neutral facial expressions (Fig. 1B), resulting in 42 trials (i.e., gaze change sequences) per block. Within a given trial, between 12 and 14 gaze changes occurred (both for active and passive viewing; Fig. 1C). The number and order of gaze changes was pseudo-randomized for each participant, both viewing conditions, and all emotional expressions. On average, 2184 gaze changes occurred per participant, corresponding to 30.3 events in each cell of the experimental design (i.e. for each combination of viewing, attended eye, emotional, and gaze change condition).

Throughout a given trial, the emotional expression of the stimulus face remained constant. At the beginning of each trial, a fixation cross was centrally presented for 1 s before the face appeared. As soon as the sequence of gaze events was completed, a screen was shown with the number of gaze changes in this trial and a second number that was higher or lower by 1. Participants had to select the correct number of gaze events by pressing one of two response buttons.

Prior to each active or passive viewing block, two practice trials were conducted. After every 10^th^ trial and between blocks, participants could rest. Throughout experimental trials, gaze position was monitored to check compliance with fixation instructions.

The experiment was programmed and controlled using *Presentation* software (version 18.10, Neurobehavioral Systems Inc, Albany, CA).

### 2.4 Data Acquisition

EEG data were recorded from 48 Ag/AgCl electrodes at a sampling rate of 500 Hz using BrainAmp amplifiers (BrainProducts GmbH, Gilching, Germany). Electrodes were mounted in an elastic cap (EasyCap, Herrsching, Germany) placed according to the international 10-10 system at positions FP1, FPz, FP2, AF7, AFz, AF8, F7, F5, F3, Fz, F4, F6, F8, FT9, FT7, FC5, FC1, FC2, FC6, FT8, FT10, T7, C3, Cz, C4, T8, CP5, CP1, CP2, CP6, P9, P7, P5, P3, Pz, P4, P6, P8, P10, PO9, PO7, POz, PO8, PO10, O1, Oz, O2, and Iz. Four additional Ag/AgCl electrodes were placed at the outer canthus and infraorbital ridge of each eye to record the electrooculogram. During recordings, the EEG signal was referenced to an electrode on the left mastoid process (A1); FCz was used as ground electrode location. An additional electrode on the right mastoid (A2) was included to ensure a symmetric electrode setup for later re-calculation to average reference. All impedances were kept below 5 kΩ.

Gaze position was recorded at a sampling rate of 500 Hz with a video-based eye tracker (iView X Hi-Speed 1250, Sensomotoric Instruments GmbH, Teltow, Germany). The eye tracking system was calibrated using a nine-point calibration grid before the experiment, before every tenth trial, in breaks between blocks or in-between trials, if necessary. Measurement error was assessed with a four-point validation procedure and kept below 0.5° for both horizontal and vertical gaze positions.

### 2.5 Data Analysis

All data processing was done in MATLAB (version 2014a, The MathWorks Inc., Natick, MA). Offline, the EEG data were re-calculated to an average reference montage. The data were then band-pass filtered using EEGLAB’s default finite response filter with the low cutoff (- 6 dB attenuation) set to 0.1 Hz and the high cutoff set to 45 Hz. Subsequently, the EEG was synchronized with the eye tracking data using the EYE-EEG (Dimigen et al., 2011) toolbox for EEGLAB (version 14.1.1; Delorme & Makeig, 2004). Eye movement artifacts in the EEG were corrected with the Multiple Source Eye Correction procedure (MSEC; Berg & Scherg, 1994; Ille, Berg, & Scherg, 2002) as implemented in the BESA software (version 6.0, BESA GmbH, Gräfeling, Germany).

The artifact-corrected continuous EEG was segmented into epochs from −700 to 1000 ms around each gaze change event and baseline-corrected with a −100 to 0 ms pre-stimulus baseline. A total of 4.2% of all epochs was excluded because voltages in at least one channel exceeded ±100 μV or because the participant’s gaze had remained in the eye region of the picture for less than 500 ms after a gaze change of the stimulus face.

To examine ERP effects of eye contact compared to gaze aversion, ERPs of the gaze conditions LD and RD, as well as DL and DR, were pooled, resulting in two gaze change categories: direct-to-averted gaze (DAv) and averted-to-direct gaze (AvD). In the following, these categories are considered as levels of a two-level factor *eye contact*.

N170 amplitudes were measured in the averaged ERPs for every participant and condition at electrode PO8 by peak detection within a time window of 130-220 ms after stimulus gaze change onset. Additionally, effects of eye contact were investigated in later ERP components. For the EPN, an electrode cluster consisting of P7, P8, PO9, PO7, PO8, PO10, O1, Oz, O2, and Iz was selected, and the mean amplitude was examined in four consecutive non-overlapping 50 ms time windows between 200 and 400 ms post-stimulus. The P300 component was quantified by peak detection at electrode Cz between 150 ms and 400 ms post-stimulus.

In addition, we investigated the brain activity related to the end of the saccade, which triggered the change of the face picture in the active condition. EEG potentials time-locked to saccade offsets are called fixation-related potentials (FRP; e.g., Yagi, 1979; Dimigen et al., 2011). Here, the eye tracker registered the arrival of the participant’s gaze in the eye region of the face (i.e. the crossing of the invisible boundary around the eye region), which precedes the actual fixation onset by a few milliseconds. Therefore, we refer to this signal as a *quasi-FRP*.

Since the picture changes only occurred 500 ms after the saccade to the eye region, the quasi-FRP signal was also contained in our data epochs that were segmented from −700 to 1000 ms around the stimulus change. The same segmentation as for the change events was done for non-change events in which no picture change followed the participant’s saccade (10% of all trials). Because change epochs contained both picture change-as well as fixation-related activity, whereas non-change epochs contained pure fixation-related activity, we were able to remove the fixation-related activity from the change epochs. Thus, non-change epochs were averaged separately for the levels of the factors *eye contact* and *attended eye* but collapsed over all emotional facial expressions to obtain FRPs with a sufficiently high signal-to-noise ratio. These average FRPs were subtracted from every change epoch of the corresponding experimental condition. These corrected change epochs provided us with the ERP of pure picture change-related activity. To ensure comparability between viewing conditions, this correction was also performed for the passive condition, subtracting non-change epochs from change epochs.

### 2.6 Statistical Analysis

Amplitude measures of N170, EPN, and P300 components were submitted to analyses of variance (ANOVA) with repeated measures on factors *viewing condition* (active, passive), *attended eye* (left, right), *emotional expression* (happy, angry, neutral), and *eye contact* (DAv, AvD). For the N170, these ANOVAs were also computed separately for active and passive viewing. N170 amplitudes after correction for overlapping FRPs in the active condition, were compared to the passive condition by a repeated-measurement ANOVA with factors *viewing condition* and *eye contact*. Again, the effect of *eye contact* was examined in separate ANOVAs for active and passive viewing.

To test whether the absolute N170 amplitude and/or eye contact effect changed over the gaze change sequence within a trial, the N170 peak amplitude was regressed on the temporal position of the stimulus within a sequence (denoted as *within-trial position*), and on the dichotomous factor *eye contact*, using multiple regression. Please note that *within-trial position* was examined only up to the 13^th^ gaze event since the 14^th^ event occurred only in 1/3 of all trials and hence did not provide enough epochs for obtaining clean ERPs.

Behavioral results of the gaze change counting task were analyzed using a repeated-measurement ANOVA with accuracy (proportion of trials with correctly counted gaze changes) as a dependent variable and *viewing condition, attended eye*, and *emotion* as predictors.

For all statistical analyses, the significance level was set to *p* < .05. The effect of eye contact was tested one-sided since we expected larger amplitudes for gaze aversion as compared to eye contact. The sphericity assumption was assessed using Mauchly’s test and adjustments were made applying Huynh-Feldt correction, if needed. Pairwise comparisons were performed between emotional categories, adapting *p* values according to the Bonferroni method. Statistical analyses were conducted in R Studio (version 1.0.143; RStudio Team, Boston, MA).

## 3 Results

### 3.1 Behavioral results

Participants performed the gaze change counting task with an accuracy of 83.0% in the active, and 89.1% in the passive condition, a difference that was significant in a repeated-measurement ANOVA, *F*(1, 20) = 8.94, *p* = .007, *η*^2^ = .058. Task performance did not vary as a function of the attended eye or the emotional face expressions.

### 3.2 N170

Figures 2A and 2B show the main effects of eye contact on the ERPs in the active and passive conditions; the N170 amplitude appears to be larger for gaze aversions (direct to averted gaze) than for eye contact (averted to direct gaze). Main effects of emotion are shown in Figures 2C and 2D, indicating smaller N170 amplitude for angry as compared to neutral and happy expressions. Notably, the entire ERP waveform in the active condition appeared to be shifted towards more negative values than in the passive condition, although its overall morphology, such as peak latencies and amplitudes, seemed to be rather similar between the conditions.

**Figure 2.**
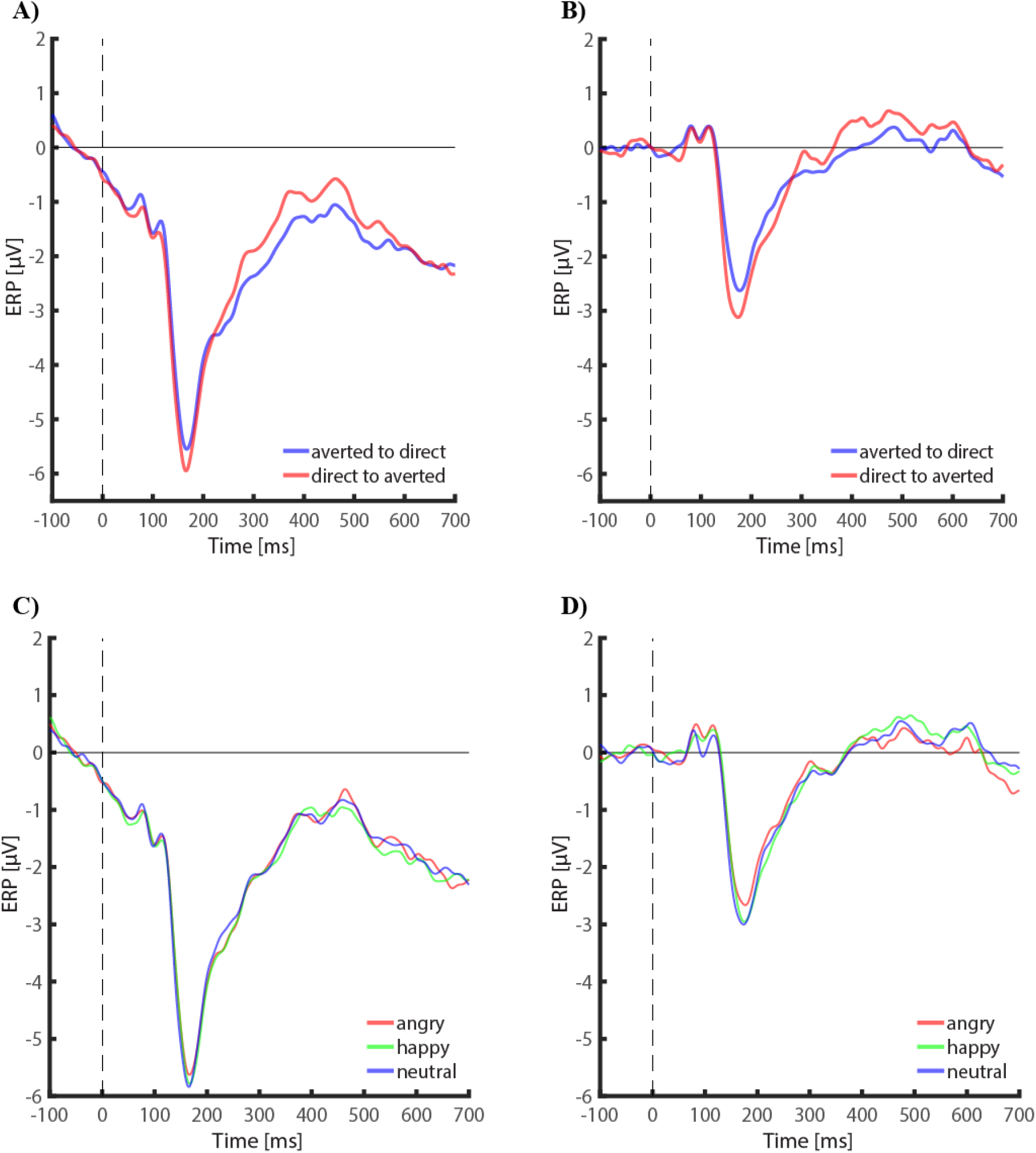
ERPs at electrode PO8 depicting the eye contact effect in the active (panel A) and passive (panel B) condition, as well as the effect of emotional face expression in the active (panel C) and passive (panel D) condition.

The time course of the topographic distribution of the eye contact effect (DA minus AD) is shown in Figure 3 for the active and passive conditions, respectively. Topographies show a focal eye contact effect around 170 ms post-stimulus at a right posterior site in both active and passive conditions.

**Figure 3.**
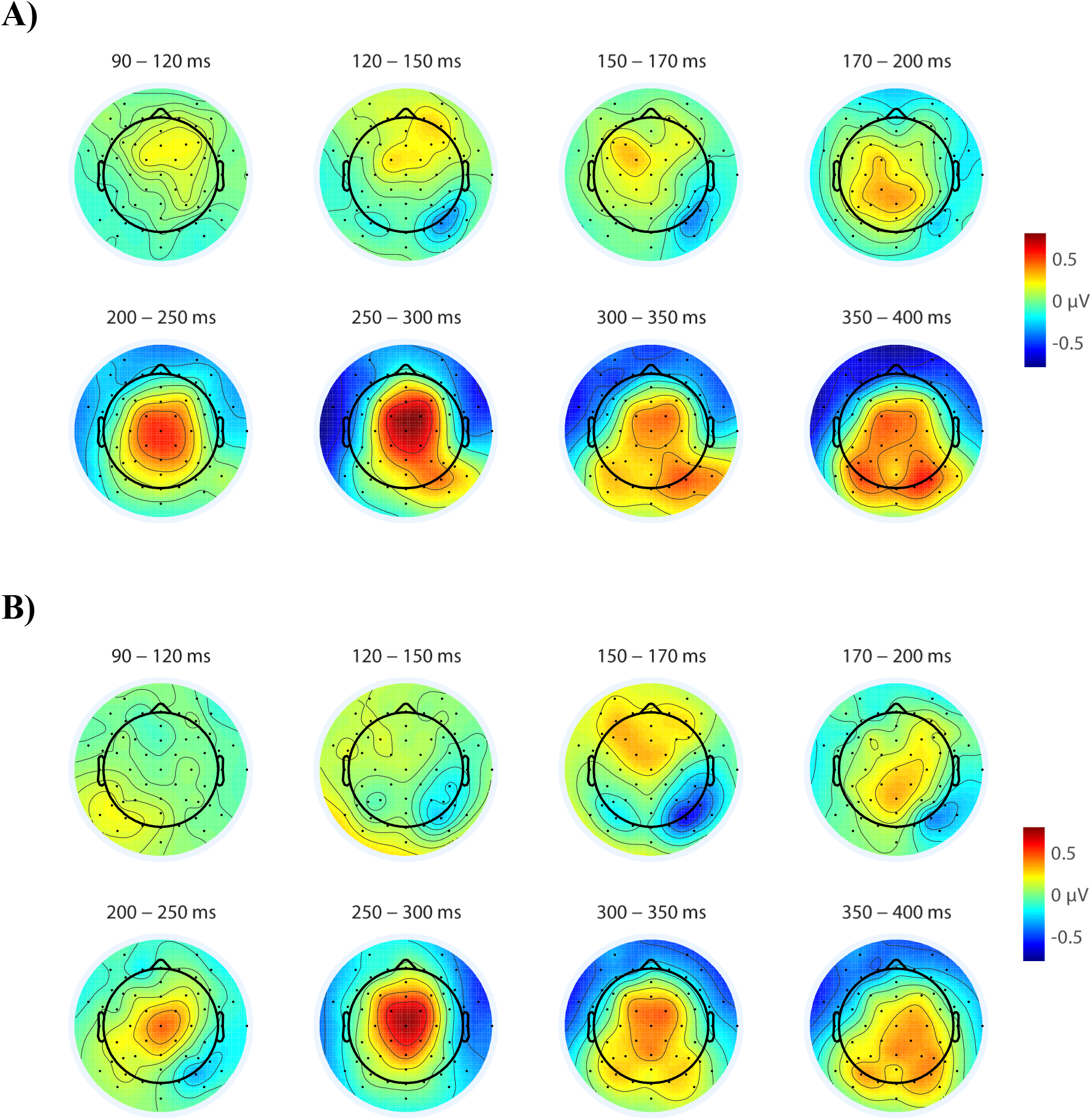
Scalp topographies of the eye contact effect (i.e., difference topographies of direct-to-averted minus averted-to-direct gaze) averaged in the indicated time windows. A) Active condition. B) Passive condition.

In the overall ANOVA of N170 amplitude at electrode PO8, main effects were found for *viewing condition*, *F*(1, 20) = 141.88, *p* < .001, η^2^ = .199, *emotion*, *F*(2, 40) = 4.81, *p* = .013, *η*^2^ = .002, and *eye contact*, *F*(1, 20) = 4.21, *p*_one-tailed_ = .027, *η*^2^ = .004 (the latter was tested one-sided according to our directed hypothesis). Furthermore, a four-way interaction of *viewing condition, attended eye, emotion*, and *eye contact* was found, *F*(2, 33.70) = 3.96, *p* = .035, *η*^2^ = .002 (for further examination see Supplement 2). No other main effects or interactions reached significance. As depicted in Figures 2C and 2D, post-hoc pairwise comparisons of emotion categories revealed significant differences between angry and happy faces, *F*(1, 20) = 8.66, *p*_adjusted_ = .024, a marginal difference between angry and neutral faces, *F*(1, 20) = 6.13, *p*_adjusted_ = .067, and no difference between happy and neutral faces.

With respect to the difference in the general appearance of the ERPs between active and passive viewing (Fig. 2), effects on the N170 were also examined in separate ANOVAs for the two viewing conditions. In passive viewing, the factor eye contact was significant, *F*(1, 20) = 5.71, *p*_one-tailed_ = .013, *η*^2^ = .008, whereas in active viewing it was only a trend, *F*(1, 20) = 2.06, *p*_one-tailed_ = .083, *η*^2^ = .002. No other effects were significant in either viewing condition.

Figure 4 illustrates the consistency of N170 amplitudes over the gaze change sequences. As tested by multiple regression, the N170 amplitude was neither modulated by *within-trial position, β* = −.001, *t*(542) = .023, *p* = .982, nor by the interaction of *within-trial position* and *eye contact, β* = .000, *t*(542) = .000, *p* = 1.000.

**Figure 4.**
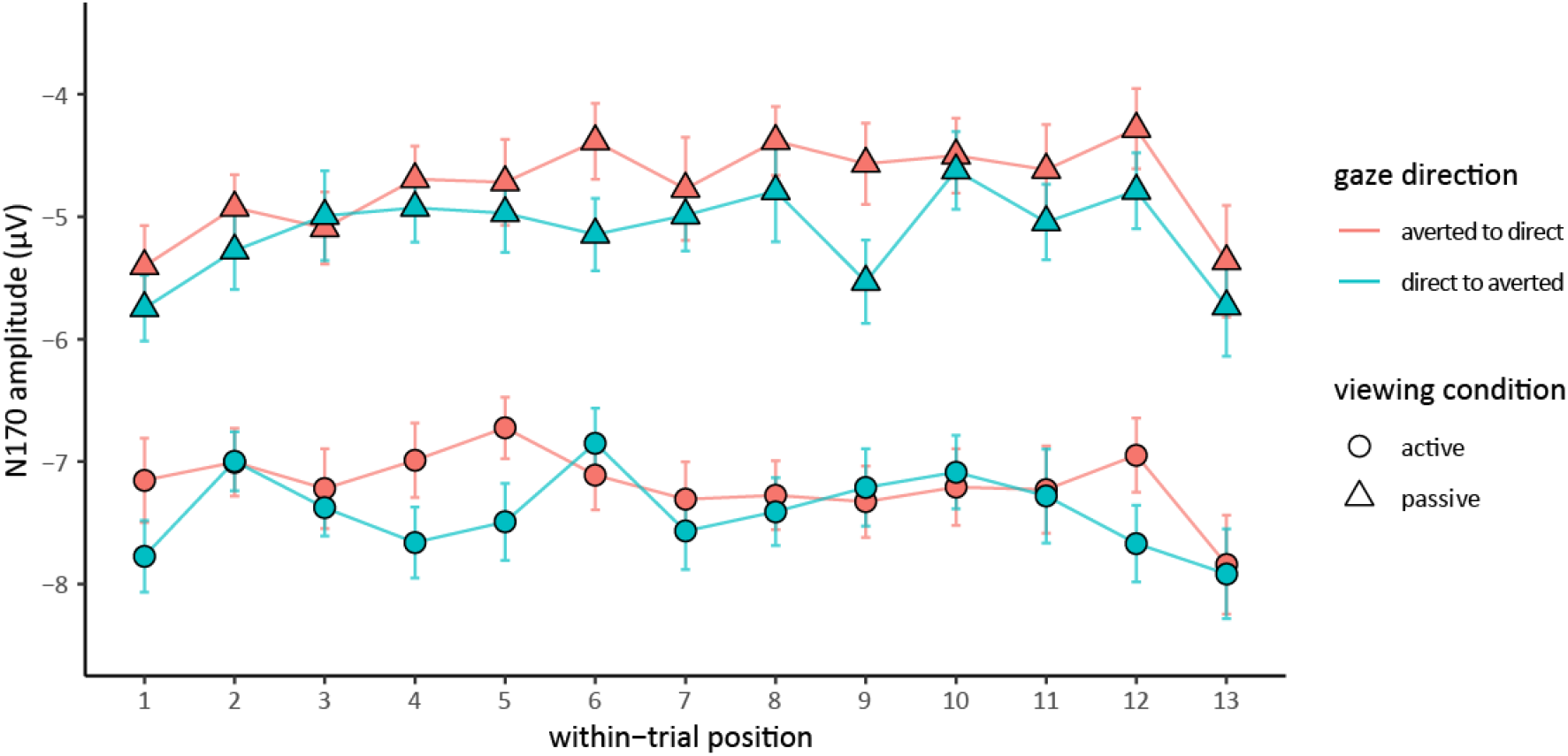
N170 amplitudes at given temporal positions within gaze change sequences for both viewing and gaze conditions at electrode PO8. Both absolute N170 amplitude as well as the N170 eye contact effect are consistent over gaze change sequences. Note that non-change events are not included in this illustration.

### 3.3 FRPs and corrected ERPs

The average fixation-related potentials (FRPs) produced by the saccade toward the eye region of the stimulus face in the active condition are shown in Figures 5A and 5B. Both in change and non-change events, a clear positive peak, the so-called lambda response, was present between 100 and 200 ms post-fixation. This peak, which is the equivalent of the visually-evoked P1 component, was followed by a small negative potential (presumably the N170 equivalent) as well as one more positive deflection (potentially a P3b component; Fig. 5A and 5B). Thereafter, the FRPs in non-change events showed a long-lasting negative deflection extending to long after the time point where gaze changes would occur in change trials (Fig. 5A); presumably, this activity was present also during gaze change events, overlapping the change-elicited ERPs (Fig. 5B). Indeed, after correcting for this overlapping activity in the active viewing condition, the ERPs in the active viewing conditions became much more similar to the passive condition (Fig. 5C and 5D).

**Figure 5.**
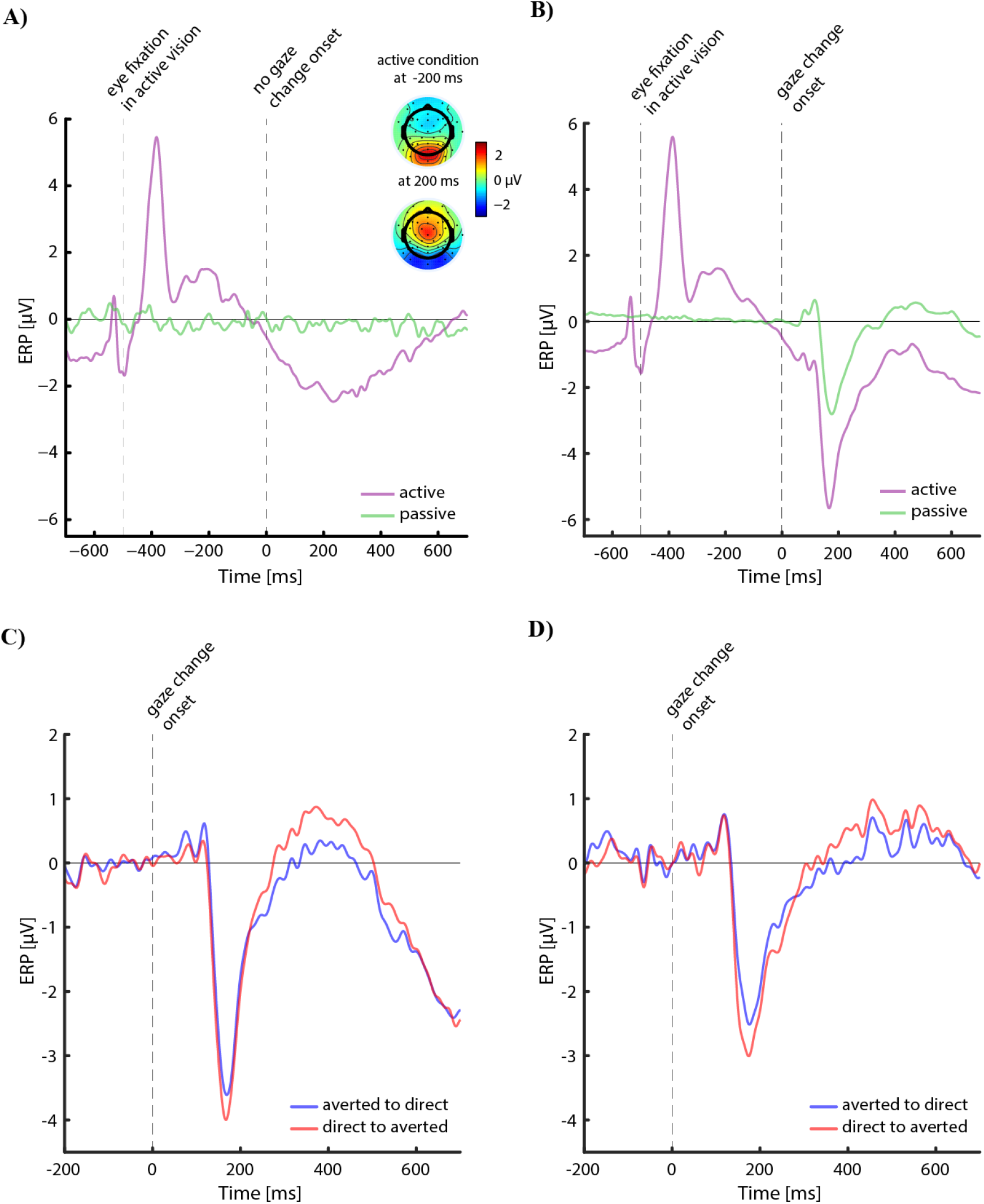
FRPs and corrected ERPs at electrode PO8. A) FRPs in non-change events. Topographies are depicted for the active condition at −200 and 200 ms. B) FRPs in change events. C) Corrected ERPs in the active condition. D) Corrected ERPs in the passive condition.

To test whether the (uncorrected) ERP differences between active and passive viewing were due to the superposition of the ERP with the preceding FRP, N170 peak amplitudes were analyzed in the corrected data. Now, the factor *viewing condition* was no longer significant, *F*(1, 20) = 1.46, *p* = .241, *η*^2^ = .007, but the factor *eye contact* still had a marked influence on (corrected) N170 amplitudes, *F*(1, 20) = 5.83, *p* = .025, *η*^2^ = .006. This held true also for separate analyses of active and passive viewing, *F*(1, 20) = 3.58, *p*_one-tailed_ = .036, *η*^2^ = .004, and *F* (1, 20) = 5.97, *p*_one-tailed_ = .024, *η*^2^ = .009, respectively.

### 3.4 EPN

The face’s emotional expression did not influence the EPN in any of the four tested time windows between 200 and 400 ms post-stimulus, *Fs*(2,40) ≤ 1.05, *ps* > .4. Analyses of the EPN time windows did not yield any other effects besides a main effect of *viewing condition*, which was presumably due to the general negative shift of the ERPs in the active condition discussed above (see also Supplement 3).

### 3.5 P300

Examination of the scalp topographies of the eye contact effect in Figure 3 suggested a central positivity starting shortly after 200 ms, as can be seen in the ERP in Figure 6. In fact, the statistical analysis of this positivity revealed that it was larger for averted gaze than for eye contact, *F*(1, 20) = 17.80, *p* < .001, *η*^2^ = .019. In addition, this central positivity was larger in active than passive viewing, as confirmed by a main effect of viewing condition, *F*(1, 20) = 66.06, *p* < .001, *η*^2^ = .383, which was presumably due to a similar superposition of ERP and preceding FRP as was found for the N170 (see *3.3 FRPs and corrected ERPs*).

**Figure 6.**
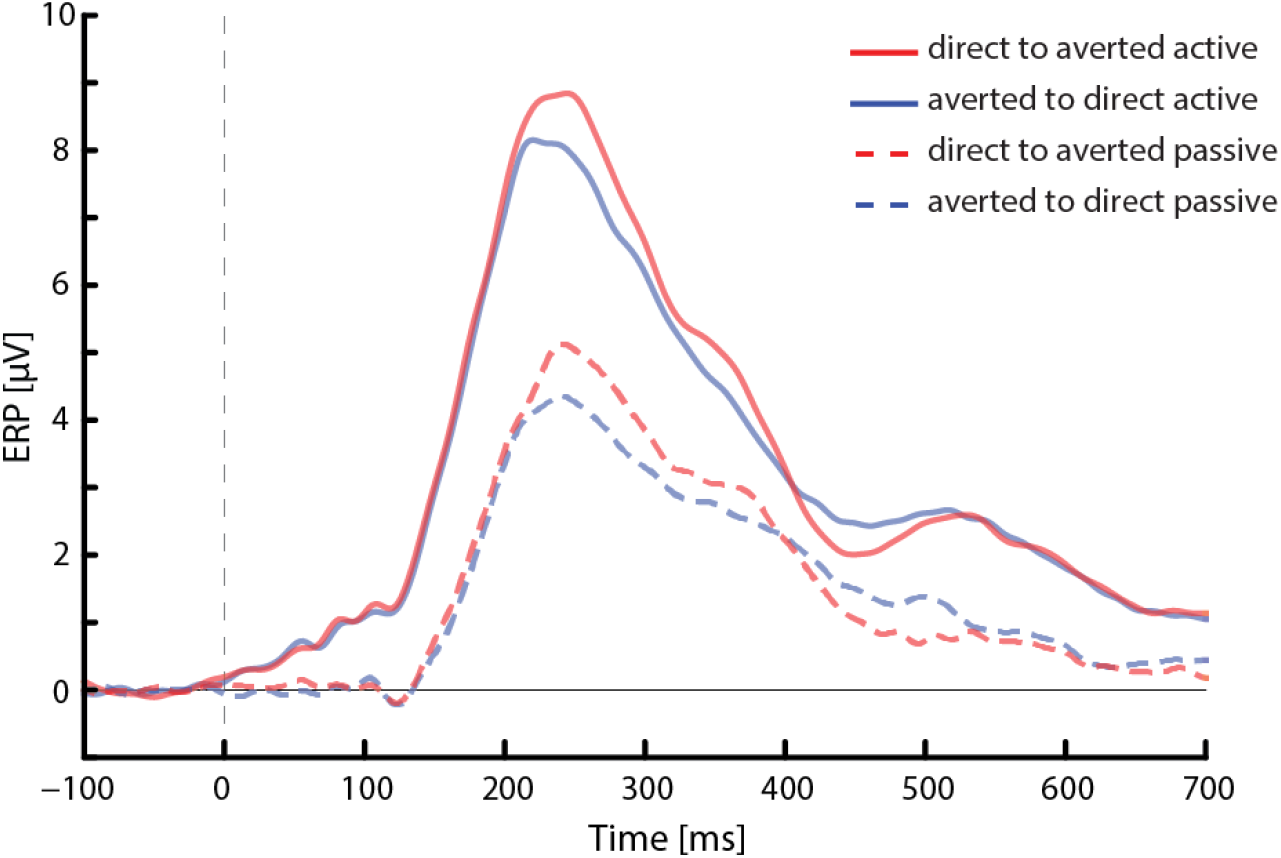
P300 effect at electrode Cz for both viewing and gaze conditions.

## 4 Discussion

The central aim of the present study was to investigate the neural correlates of eye contact in settings that more closely resemble naturalistic interactions. In a continuous-trial design, participants viewed dynamic gaze changes of faces with a happy, angry, or neutral expression. The passive observation of gaze changes was compared to an interactive condition in which gaze shifts of the observed face occurred in response to the participant establishing eye contact. With continuous stimulus presentation, we were able to replicate typical eye contact effects on the N170 as they are commonly found with conventional trial-by-trial presentation of faces. Furthermore, we found that after correcting ERPs for the effects of temporally overlapping fixation-related potentials, the eye contact effects fully generalized from the passive to the active condition. In addition, we observed (1) robust increases of a late positive component in response to eye contact and (2) significant effects of emotional facial expressions on the N170 component. In the following, we will discuss these findings in more detail.

### 4.1 Effect of eye contact on N170 amplitude

Our finding of larger N170 amplitudes for gaze aversion as compared to eye contact matches the results of previous studies using non-social, spatial tasks (Bagherzadeh-Azbari et al., in preparation; Latinus et al., 2015; Puce et al., 2000). In particular, a face changing its gaze direction from direct to averted indicates an attention shift away from the observer towards the environment. This conveys important information about the communication partners current state of mind, and about his or her intentions. Puce et al. (2015) and Latinus et al. (2015) suggested two distinct processing modes for this information, a *socially aware mode*, which is activated when social cues need to be attended and a *non-social default mode*. A social task (e.g., detection of eye contact) may activate the *socially aware mode*, which then enhances the N170 amplitude over the right hemisphere also in response to direct gaze. In the current study, however, participants counted gaze changes, which is a non-social task. Therefore, the participants’ gaze perception system was presumably in the *non-social default mode*, associated with larger N170 amplitudes in response to gaze aversion as compared to eye contact. Importantly, the gaze interaction with the stimulus in the active condition did not seem to change the *non-social default mode* to the *socially aware mode* as no interaction effect of viewing condition and gaze direction emerged.

#### 4.1.1 Continuous stimulus presentation

So far, eye contact effects in the N170 have mostly been shown for gaze changes of faces that were only briefly presented in a trial-by-trial fashion (e.g, Conty et al., 2007; Conty et al., 2012; Latinus et al., 2015; Pönkänen et al., 2011; but see Puce et al., 2000 for a continuous paradigm). In contrast, the current study investigated the eye contact effect for faces that remained on the screen and changed their gaze direction up to 13 times within a given trial. Importantly, we found that neither the absolute N170 amplitude nor the size of the N170 eye contact effect decreased over these long sequences of gaze changes (Fig. 4). If there is no attenuation of the eye contact effect over 13 successive gaze change events, it is reasonable to assume that this does also not occur for even longer sequences, such as in real-world dyadic interactions. Hence, the N170 component appears to be a promising target for further investigation of the eye contact effect in more naturalistic settings.

#### 4.1.2 Generalizing the eye contact effect to interactive viewing

In order to take a second step towards studying gaze perception under more naturalistic conditions, we recorded the N170 eye contact effect also in a gaze-contingent presentation paradigm. Specifically, the stimulus face changed its gaze direction shortly after the participant looked at the eye region, as it commonly takes place also in dyadic interactions. Interestingly, the main effect of gaze direction in the overall ANOVA suggests that the N170 eye contact effect is independent of active or passive viewing conditions. Furthermore, at least at a descriptive level, the eye contact effect was present in both viewing conditions (Fig. 2A and 2B) with comparable scalp topographies (Fig. 3). However, when computing separate ANOVAs for active and passive viewing, the eye contact effect in active viewing alone did not reach significance. As the entire ERP in the active condition was shifted towards more negative values as compared to the passive condition (Fig. 2A and 2B), we assume the extensive eye movements in the active condition to be responsible for the attenuated eye contact effect.

It is well-known that in addition to eye movement artifacts – which were successfully removed here – each saccade offset is followed by a sequence of visually-evoked brain responses, so called fixation-related potentials (FRPs; Dimigen et al., 2011; Yagi, 1979). The morphology of the FRP when participants moved their gaze to the eye region of the picture (Fig. 5A and 5B) resembled the one observed in tasks like reading (e.g., Dimigen et al., 2011; Dimigen, Kliegl, & Sommer, 2012) or face perception (Buonocore, Dimigen, & Melcher, 2019; Mares, Smith, Johnson, & Senju, 2016; Soto et al., 2018). In addition, a slow negative deflection was observed peaking at around 700 ms after eye fixation, that is, 200 ms after the stimulus gaze change in change trials (Fig. 5A). This negative deflection therefore overlapped with the gaze-change ERP (Fig. 5B), which led to a more negative curvature of the ERP in the active as compared to the passive condition.

To support this notion, we corrected gaze-change ERPs by subtracting EEG activity of corresponding non-change events to obtain a pure gaze-change-related signal (Fig. 5C and 5D). In fact, the viewing condition effect on the N170 amplitude disappeared in this analysis, which implies that there is no fundamental difference in the N170 component whether a gaze change is perceived passively or is elicited actively. Even more notably, after subtracting the overlapping FRPs, the ERPs in the active condition now also revealed a significant main effect of eye contact, indicating that this effect was indeed obscured by the overlapping activity.

However, these findings should be treated with some caution since non-change events occurred more rarely than change events (proportion of 0.1 versus 0.9). Thus, the negative deflection at around 200 ms after expected picture changes in non-change events might reflect a violation of the change expectation, which is also supported by its topography, showing a typical P3a component pattern (Fig. 5A). However, this potential was only present in the active condition although picture changes occurred with the same probability in the passive condition. To further examine this issue, future studies involving gaze-contingent stimuli could vary the proportion of change and non-change events systematically. Another possibility to disentangle the overlapping brain potentials from saccades and gaze changes in this paradigm would be to jitter the interval between the saccade and the resulting gaze change. This would allow using modern regression-based analysis techniques (Ehinger & Dimigen, 2018; Smith & Kutas, 2015) to disentangle both types of potentials.

Taken all together, the N170 eye contact effect seems indeed to be generalizable to an interactive viewing setting, when considering the overlap with preceding FRPs.

### 4.2 Interactions between eye contact and emotional facial expressions

We investigated whether gaze direction affects the processing of emotional facial expressions reflected in the EPN, and vice versa, whether emotional expressions influenced the N170 eye contact effect. Yet, we did not find any evidence for an interaction of gaze direction and facial expression, neither in the EPN nor in the N170. This stands in contrast to a recent study (Bagherzadeh-Azbari et al., in preparation) in which typical EPN effects were found for gaze changes from averted to direct (eye contact) but not for direct to averted (gaze aversion). Crucially, this study used the same stimuli as the present study but presented them trial-by-trial, that is, such that each gaze event was separated by an inter-trial blank screen and a behavioral response. The absence of EPN effects in both our active and passive condition thus indicates that such effects might be restricted to conventional trial-wise picture presentation and might not generalize to more naturalistic conditions in which the same face remains continuously visible.

Although we observed no interaction, the emotional expression of the face did influence N170 amplitude as a main effect. Specifically, angry faces elicited smaller amplitudes than happy and neutral faces (Fig. 2C and 2D). This finding was unexpected since the N170 is not thought to be involved in processing emotional content (Eimer, 2011; Rellecke, Sommer, & Schacht, 2013). However, given that the iris-pupil complex and its contrast to the white sclera play an important role in generating the N170 (Puce et al., 2015; Rossi, Parada, Kolchinsky, & Puce, 2014), N170 amplitude differences between emotional categories might have been due to differences in the amount of visible white sclera between emotional categories (happy > angry > neutral; see Supplement 4 for measurements and statistics). Future studies might systematically investigate the effect of visible eye size on the N170 irrespective of gaze direction and emotional expression.

### 4.3 Eye contact effect on the P300 component

In the scalp topographies of the eye contact effects, we observed strong positive potentials at central electrodes from about 200 ms onwards, in both the active and passive conditions (Fig. 3 and 6). This pattern of a fronto-central positivity followed by a parieto-central positivity resembles a P3a component followed by a P3b component. As statistically confirmed, P300 amplitudes were larger for gaze aversion than for eye contact. Since the P300 can be interpreted as an index of attentional processing (Cuthbert et al., 2000; Donchin & Coles, 1988), these findings suggest that gaze changes from direct to averted gaze draw more attention than gaze changes from averted to direct, which stands in contrast to previous research. Thus, Conty et al. (2007) and Myllyneva and Hietanen (2015), reported P300 effects in the opposite direction, that is, larger amplitudes for eye contact than for gaze aversion. Furthermore, paying more attention to averted than to direct gaze would contradict the theory that eye contact rather than gaze aversion increases the saliency of a face. Yet, our findings are consistent with a previously reported P350 component which was larger for averted than for direct gaze (Puce et al., 2000). Future research should clarify the conditions under which eye contact elicits larger or smaller P300 responses than averted gaze. Relevant modulating factors for this could be the task and the relative probabilities of the critical events.

### 4.4 Conclusions and perspectives

The present findings show that eye contact effects in the N170 occur in a similar way, regardless of whether they are elicited during passive viewing or in a more interactive situation. Furthermore, we show that these N170 effects are not attenuated across a long sequence of gaze changes made by the same face. However, we could not replicate emotion effects on the EPN component, which questions whether these effects generalize to continuous-trial designs. We consider these findings as relevant steps towards investigating dyadic interactions in more natural settings. Nevertheless, it should be emphasized that the current interactive viewing paradigm is still a strong simplification of the complexities of real-world gaze interactions. For example, the observed stimuli did not take actions on their own but merely responded to the actions of the participant. Hence, one important next step would be to implement a paradigm in which eye contact is initiated by the participant and stimulus face alternately to study truly mutual interactions, before eventually being able to move on to real dyadic interactions between two humans.

## Acknowledgements

We wish to thank Thomas Pinkpank and Rainer Kniesche for technical support, Ulrike Bunzenthal for assistance with the EEG recordings, and Meryem Giden for help with the preparation of the stimulus material.

## Open Practice Statement

All stimulus material, analysis scripts, and preprocessed EEG data (in compliance with the current European General Data Protection Regulation) are available from the authors upon reasonable request. The experiment was not preregistered.

## References

Bagherzadeh-Azbari, S., Lion, C. J., Dimigen, O., Stephani, T., & Sommer, W. (in preparation). The Interplay of Eye Gaze and Emotional Facial Expressions: An Event-Related Brain Potential Study.

Bentin, S., Allison, T., Puce, A., Perez, E., & McCarthy, G. (1996). Electrophysiological Studies of Face Perception in Humans. Journal of Cognitive Neuroscience, 8(6), 551–565. https://doi.org/10.1162/jocn.1996.8.6.551

Berg, P., & Scherg, M. (1994). A multiple source approach to the correction of eye artifacts. Electroencephalography and Clinical Neurophysiology, 90(3), 229–241. https://doi.org/10.1016/0013-4694(94)90094-9

Bublatzky, F., Pittig, A., Schupp, H. T., & Alpers, G. W. (2017). Face-to-face: Perceived personal relevance amplifies face processing. Social Cognitive and Affective Neuroscience, 12(5), 811–822. https://doi.org/10.1093/scan/nsx001

Buonocore, A., Dimigen, O., & Melcher, D. (2019). Post-saccadic face processing is modulated by pre-saccadic preview: Evidence from fixation-related potentials. BioRxiv. Advance online publication. https://doi.org/10.1101/610717

Conty, L., Coelho, E., Tijus, C., Hugueville, L., & George, N. (2006). Searching for asymmetries in the detection of gaze contact versus averted gaze under different head views: A behavioural study. Spatial Vision, 19(6), 529–545. https://doi.org/10.1163/156856806779194026

Conty, L., Dezecache, G., Hugueville, L., & Grezes, J. (2012). Early binding of gaze, gesture, and emotion: neural time course and correlates. The Journal of Neuroscience: the Official Journal of the Society for Neuroscience, 32(13), 4531–4539. https://doi.org/10.1523/JNEUROSCI.5636-11.2012

Conty, L., N’Diaye, K., Tijus, C., & George, N. (2007). When eye creates the contact! ERP evidence for early dissociation between direct and averted gaze motion processing. Neuropsychologia, 45(13), 3024–3037. https://doi.org/10.1016/j.neuropsychologia.2007.05.017

Cuthbert, B. N., Schupp, H. T., Bradley, M. M., Birbaumer, N., & Lang, P. J. (2000). Brain potentials in affective picture processing: Covariation with autonomic arousal and affective report. Biological Psychology, 52(2), 95–111. https://doi.org/10.1016/S0301-0511(99)00044-7

Delorme, A., & Makeig, S. (2004). Eeglab: An open source toolbox for analysis of single-trial EEG dynamics including independent component analysis. Journal of Neuroscience Methods, 134(1), 9–21. https://doi.org/10.1016/j.jneumeth.2003.10.009

Dimigen, O., Kliegl, R., & Sommer, W. (2012). Trans-saccadic parafoveal preview benefits in fluent reading: A study with fixation-related brain potentials. NeuroImage, 62(1), 381–393. https://doi.org/10.1016/j.neuroimage.2012.04.006

Dimigen, O., Sommer, W., Hohlfeld, A., Jacobs, A. M., & Kliegl, R. (2011). Coregistration of eye movements and EEG in natural reading: Analyses and review. Journal of Experimental Psychology. General, 140(4), 552–572. https://doi.org/10.1037/a0023885

Donchin, E., & Coles, M. G. H. (1988). Is the P300 component a manifestation of context updating? The Behavioral and Brain Sciences, 11(03), 357. https://doi.org/10.1017/S0140525X00058027

Ehinger, B. V., & Dimigen, O. (2018). Unfold: An integrated toolbox for overlap correction, non-linear modeling, and regression-based EEG analysis. BioRxiv. Advance online publication. https://doi.org/10.1101/360156

Eimer, M. (2000). The face-specific N170 component reflects late stages in the structural encoding of faces. Neuroreport, 11(10), 2319–2324.

Eimer, M. (2011). The face-sensitive N170 component of the event-related brain potential. In A. J. Calder (Ed.), Oxford library of psychology. The Oxford handbook of face perception (1st ed., pp. 329–344). Oxford: Oxford Univ. Press.

Hamilton, A. F. d. C. (2016). Gazing at me: the importance of social meaning in understanding direct-gaze cues. Philosophical Transactions of the Royal Society of London. Series B, Biological Sciences, 371(1686), 20150080. https://doi.org/10.1098/rstb.2015.0080

Hietanen, J. K., Leppanen, J. M., Peltola, M. J., Linna-Aho, K., & Ruuhiala, H. J. (2008). Seeing direct and averted gaze activates the approach-avoidance motivational brain systems. Neuropsychologia, 46(9), 2423–2430. https://doi.org/10.1016/j.neuropsychologia.2008.02.029

Ille, N., Berg, P., & Scherg, M. (2002). Artifact correction of the ongoing EEG using spatial filters based on artifact and brain signal topographies. Journal of Clinical Neurophysiology: Official Publication of the American Electroencephalographic Society, 19(2), 113–124.

Itier, R. J., Alain, C., Kovacevic, N., & McIntosh, A. R. (2007). Explicit versus implicit gaze processing assessed by ERPs. Brain Research, 1177, 79–89. https://doi.org/10.1016/j.brainres.2007.07.094

Kendon, A. (1967). Some functions of gaze-direction in social interaction. Acta Psychologica,26(1), 22–63.

Klucharev, V., & Sams, M. (2004). Interaction of gaze direction and facial expressions processing: ERP study. Neuroreport, 15(4), 621–625.

Langner, O., Dotsch, R., Bijlstra, G., Wigboldus, D. H. J., Hawk, S. T., & van Knippenberg, A. (2010). Presentation and validation of the Radboud Faces Database. Cognition & Emotion,24(8), 1377–1388. https://doi.org/10.1080/02699930903485076

Latinus, M., Love, S. A., Rossi, A., Parada, F. J., Huang, L., Conty, L., … Puce, A. (2015). Social decisions affect neural activity to perceived dynamic gaze. Social Cognitive and Affective Neuroscience, 10(11), 1557–1567. https://doi.org/10.1093/scan/nsv049

Lusk, L. G., & Mitchel, A. D. (2016). Differential Gaze Patterns on Eyes and Mouth During Audiovisual Speech Segmentation. Frontiers in Psychology, 7, 52. https://doi.org/10.3389/fpsyg.2016.00052

Lyyra, P., Astikainen, P., & Hietanen, J. K. (2017). Look at them and they will notice you: Distractor-independent attentional capture by direct gaze in change blindness. Visual Cognition, 1–12. https://doi.org/10.1080/13506285.2017.1370052

Mares, I., Smith, M., Johnson, M. H., & Senju, A. (2016). Direct gaze facilitates rapid orienting to faces: Evidence from express saccades and saccadic potentials. Biological Psychology,121(Pt A), 84–90. https://doi.org/10.1016/j.biopsycho.2016.10.003

Myllyneva, A., & Hietanen, J. K. (2015). There is more to eye contact than meets the eye. Cognition, 134, 100–109. https://doi.org/10.1016/j.cognition.2014.09.011

Oldfield, R. C. (1971). The assessment and analysis of handedness: The Edinburgh inventory. Neuropsychologia, 9(1), 97–113. https://doi.org/10.1016/0028-3932(71)90067-4

Pelphrey, K. A., Morris, J. P., & McCarthy, G. (2005). Neural basis of eye gaze processing deficits in autism. Brain: a Journal of Neurology, 128(Pt 5), 1038–1048. https://doi.org/10.1093/brain/awh404

Pönkänen, L. M., Alhoniemi, A., Leppanen, J. M., & Hietanen, J. K. (2011). Does it make a difference if I have an eye contact with you or with your picture? An ERP study. Social Cognitive and Affective Neuroscience, 6(4), 486–494. https://doi.org/10.1093/scan/nsq068

Puce, A., Latinus, M., Rossi, A., daSilva, E., Parada, F. J., Love, S., … Jayaraman, S. (2015). Neural Bases for Social Attention in Healthy Humans. In A. Puce & B. I. Bertenthal (Eds.), The Many Faces of Social Attention: Behavioral and Neural Measures (1st ed., pp. 93–127). Cham: Springer. https://doi.org/10.1007/978-3-319-21368-2_4

Puce, A., Smith, A., & Allison, T. (2000). Erps evoked by viewing facial movements. Cognitive Neuropsychology, 17(1), 221–239. https://doi.org/10.1080/026432900380580

Rellecke, J., Palazova, M., Sommer, W., & Schacht, A. (2011). On the automaticity of emotion processing in words and faces: event-related brain potentials evidence from a superficial task. Brain and Cognition, 77(1), 23–32. https://doi.org/10.1016/j.bandc.2011.07.001

Rellecke, J., Sommer, W., & Schacht, A. (2013). Emotion effects on the n170: A question of reference? Brain Topography, 26(1), 62–71. https://doi.org/10.1007/s10548-012-0261-y

Richardson, D. C., Dale, R., & Kirkham, N. Z. (2007). The art of conversation is coordination: Common ground and the coupling of eye movements during dialogue. Psychological Science, 18(5), 407–413. https://doi.org/10.1111/j.1467-9280.2007.01914.x

Rossi, A., Parada, F. J., Kolchinsky, A., & Puce, A. (2014). Neural correlates of apparent motion perception of impoverished facial stimuli: a comparison of ERP and ERSP activity. NeuroImage, 98, 442–459. https://doi.org/10.1016/j.neuroimage.2014.04.029

Schacht, A., & Sommer, W. (2009). Emotions in word and face processing: early and late cortical responses. Brain and Cognition, 69(3), 538–550. https://doi.org/10.1016/j.bandc.2008.11.005

Schilbach, L., Timmermans, B., Reddy, V., Costall, A., Bente, G., Schlicht, T., & Vogeley, K. (2013). Toward a second-person neuroscience. The Behavioral and Brain Sciences, 36(4), 393–414. https://doi.org/10.1017/S0140525X12000660

Schneier, F. R., Rodebaugh, T. L., Blanco, C., Lewin, H., & Liebowitz, M. R. (2011). Fear and avoidance of eye contact in social anxiety disorder. Comprehensive Psychiatry, 52(1), 81–87. https://doi.org/10.1016/j.comppsych.2010.04.006

Schupp, H. T., Junghöfer, M., Weike, A. I., & Hamm, A. O. (2003). Emotional facilitation of sensory processing in the visual cortex. Psychological Science, 14(1), 7–13. https://doi.org/10.1111/1467-9280.01411

Schupp, H. T., Ohman, A., Junghöfer, M., Weike, A. I., Stockburger, J., & Hamm, A. O. (2004). The facilitated processing of threatening faces: an ERP analysis. Emotion (Washington, D.C.), 4(2), 189–200. https://doi.org/10.1037/1528-3542.4.2.189

Senju, A., & Hasegawa, T. (2005). Direct gaze captures visuospatial attention. Visual Cognition,12(1), 127–144. https://doi.org/10.1080/13506280444000157

Senju, A., & Johnson, M. H. (2009). The eye contact effect: mechanisms and development. Trends in Cognitive Sciences, 13(3), 127–134. https://doi.org/10.1016/j.tics.2008.11.009

Smith, N., & Kutas, M. (2015). Regression-based estimation of ERP waveforms: I. The rERP framework. Psychophysiology, 52(2), 157–168. https://doi.org/10.1111/psyp.12317

Soto, V., Tyson-Carr, J., Kokmotou, K., Roberts, H., Cook, S., Fallon, N., … Stancak, A. (2018). Brain Responses to Emotional Faces in Natural Settings: A Wireless Mobile EEG Recording Study. Frontiers in Psychology, 9, 2003. https://doi.org/10.3389/fpsyg.2018.02003

Tso, I. F., Mui, M. L., Taylor, S. F., & Deldin, P. J. (2012). Eye-contact perception in schizophrenia: relationship with symptoms and socioemotional functioning. Journal of Abnormal Psychology, 121(3), 616–627. https://doi.org/10.1037/a0026596

Vertegaal, R., Slagter, R., van der Veer, G., & Nijholt, A. (2001). Eye gaze patterns in conversations. In J. Jacko (Ed.), Proceedings of the SIGCHI Conference on Human Factors in Computing Systems (pp. 301–308). New York, NY: ACM. https://doi.org/10.1145/365024.365119

Yagi, A. (1979). Saccade size and lambda complex in man. Physiological Psychology, 7(4), 370–376. https://doi.org/10.3758/BF03326658

