## Supplement for "Eye contact in active and passive viewing: event-related brain potential evidence from a combined eye tracking and EEG study"

### Supplement 1. Stimulus material: IDs of pictures taken from the Radboud Faces Database

female_01, female_04, female_18, female_26, female_57, female_58, female_61, male_03, male_15, male_21, male_23, male_24, male_36, male_71.

### Supplement 2. Further investigation of the four-way interaction in the overall N170 ANOVA

In the overall ANOVA of the N170 peak amplitude, a four-way interaction of factors viewing condition x attended eye x emotion x eye contact emerged. Post-hoc analyses showed that this effect might derive from emotion by eye contact interactions which differ between active and passive vision but only when attending the right eye. While N170 amplitudes show clear increases (i.e., negative slope in the plots) only for angry faces in the active-right condition, this is true only for happy and neutral faces in the passive-right condition (upper right panel in Figure S2).

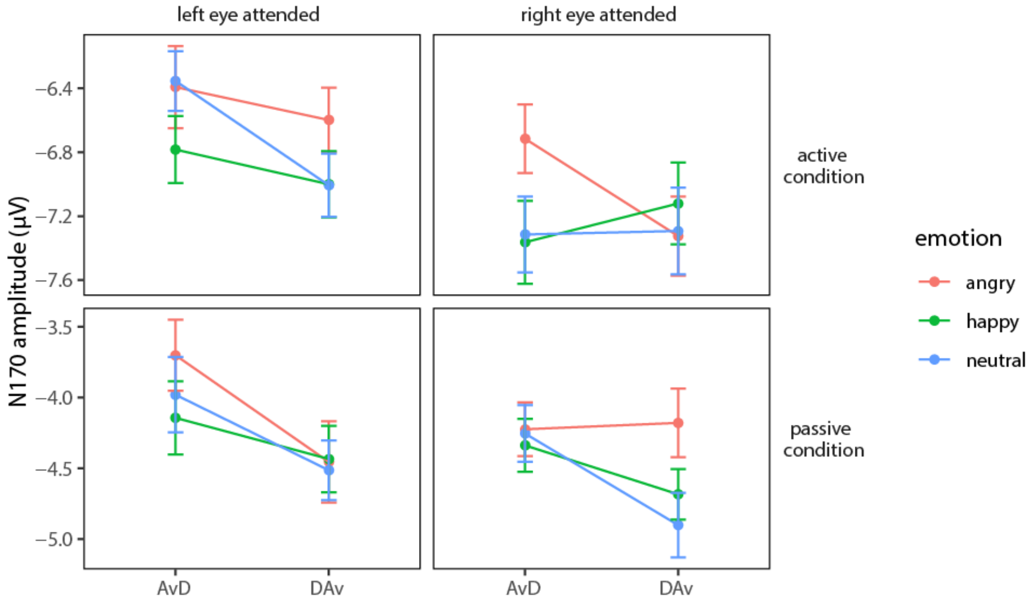


**Figure S2.** N170 peak amplitudes at PO8 for averted-to-direct (AvD) and direct-to-averted (DAv) gaze, separately depicted for active (top panels) and passive condition (bottom panels), each for left and right eye attended (left and right panels, respectively), and emotional category (color-coded). Error bars show the standard error of the mean.

**Supplement 3. Visualization of emotion effects on the EPN**

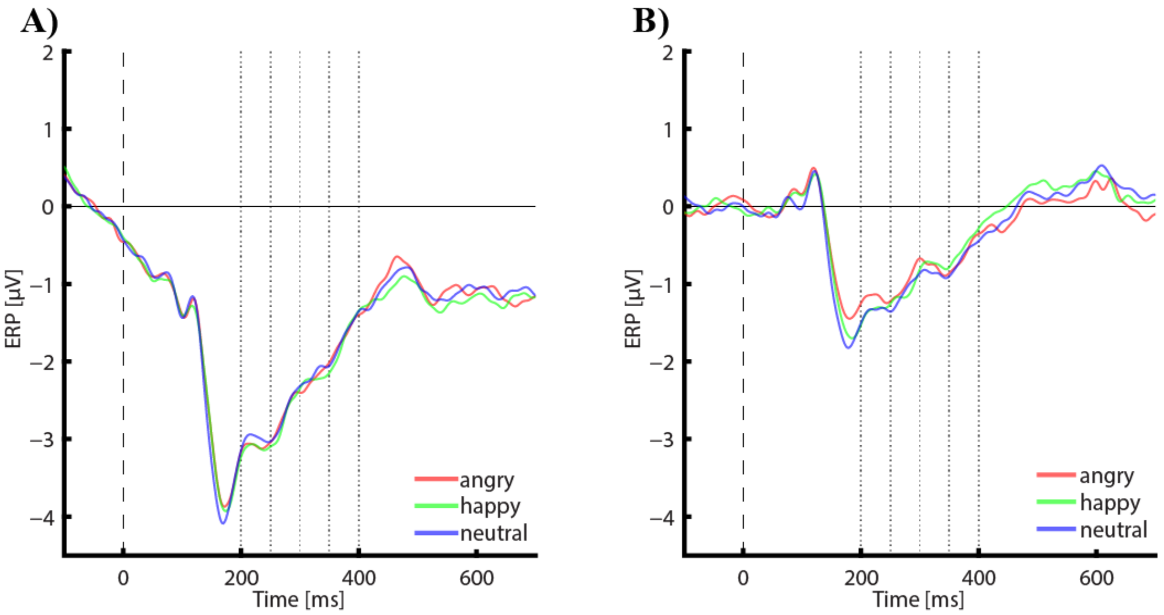


**Figure S3.** Emotion does not affect the EPN (electrodes P7/8, PO9/10, PO7/8, O1/2, Oz, Iz). A) Active condition. B) Passive condition. The dotted lines depict the boundaries of the time windows for which statistics were applied.

**Supplement 4. Eye sizes of the stimuli**

The distances between upper and lower eye lids of the stimuli were measured (in pixels) to examine differences in the amount of visible sclera between emotional conditions. The analysis revealed mean distances of 13.69, 16.30, and 17.92 pixels for happy, angry, and for neutral faces, respectively. Bonferroni-corrected pairwise-comparisons showed that all emotions were significantly different (all *p*s < .001). These differences in visible sclera could explain smaller N170 amplitudes in response to angry versus neutral faces. However, this account is inconsistent with the comparison of angry and happy faces: While happy eyes were narrower than angry eyes, N170 amplitudes were larger in happy than in angry eyes.
